# Zombi: A phylogenetic simulator of trees, genomes and sequences that accounts for dead lineages

**DOI:** 10.1101/339473

**Authors:** Adrián A. Davín, Théo Tricou, Eric Tannier, Damien M. de Vienne, Gergely J. Szöllősi

## Abstract

**Summary:** Here we present **Zombi**, a tool to simulate the evolution of species, genomes and sequences *in silico*, that considers for the first time the evolution of genomes in extinct lineages. It also incorporates various features that have not to date been combined in a single simulator, such as the possibility of generating species trees with a pre-defined variation of speciation and extinction rates through time, simulating explicitly intergenic sequences of variable length and outputting gene tree - species tree reconciliations.

**Availability and implementation:** Source code and manual are freely available in https://github.com/AADavin/ZOMBI/

## 1. Introduction

Reconstructing the pattern of horizontal gene transfers between species can help us date the origin of different taxa (Davín et al. 2018; Wolfe and Fournier 2018), understand the spread of genes of clinical importance (Lerminiaux and Cameron 2019) and resolve difficult phylogenetic questions, such inferring the rooting point of prokaryotic trees (Abby et al. 2012; Szöllősi et al. 2012; Williams et al. 2017) or the evolutionary position of certain lineages of unclear origin (Boussau, Guéguen, and Gouy 2008). In the last decades, a large number of simulators have been developed to model a wide range of evolutionary scenarios (Dalquen et al. 2011; Mallo, De Oliveira Martins, and Posada 2016; Beiko and Charlebois 2007; Sjöstrand et al. 2013; Carvajal-Rodríguez 2008; Kundu and Bansal 2019) but none so far have considered the existence of extinct lineages and the horizontal transmission of genes (by lateral gene transfers) involving species that are not represented in the phylogeny (see: (Szöllősi et al. 2013; Fournier, Huang, and Gogarten 2009; Zhaxybayeva and Peter Gogarten 2004)). Zombi simulates explicitly the genome evolution taking place in these *extinct* lineages, which is expected to have an impact in extant lineages by means of Lateral Gene Transfers (Szöllősi et al. 2013). By not considering extinct lineages, other simulators make the implicit assumption that the transfer donor always leaves a surviving descendant among sampled species, while we know that this is most often not true (Szöllősi et al. 2013). Making this assumption may potentially hamper our ability to simulate realistic scenarios of evolution. In addition to considering evolution along extinct lineages, Zombi includes several features hitherto not found together in any other simulator (Table S1).

## 2. Basic features of Zombi

Zombi is a multilevel simulator, where a species tree is first simulated, then genomes evolve along the branches of this species tree, and finally, sequences are generated for each genome. These three steps, depicted in Figure 1 and detailed hereafter, are controlled by three main “modes”, named **T, G**, and **S**, for species **T**ree, **G**enome and **S**equence, respectively.

**Fig 1.**
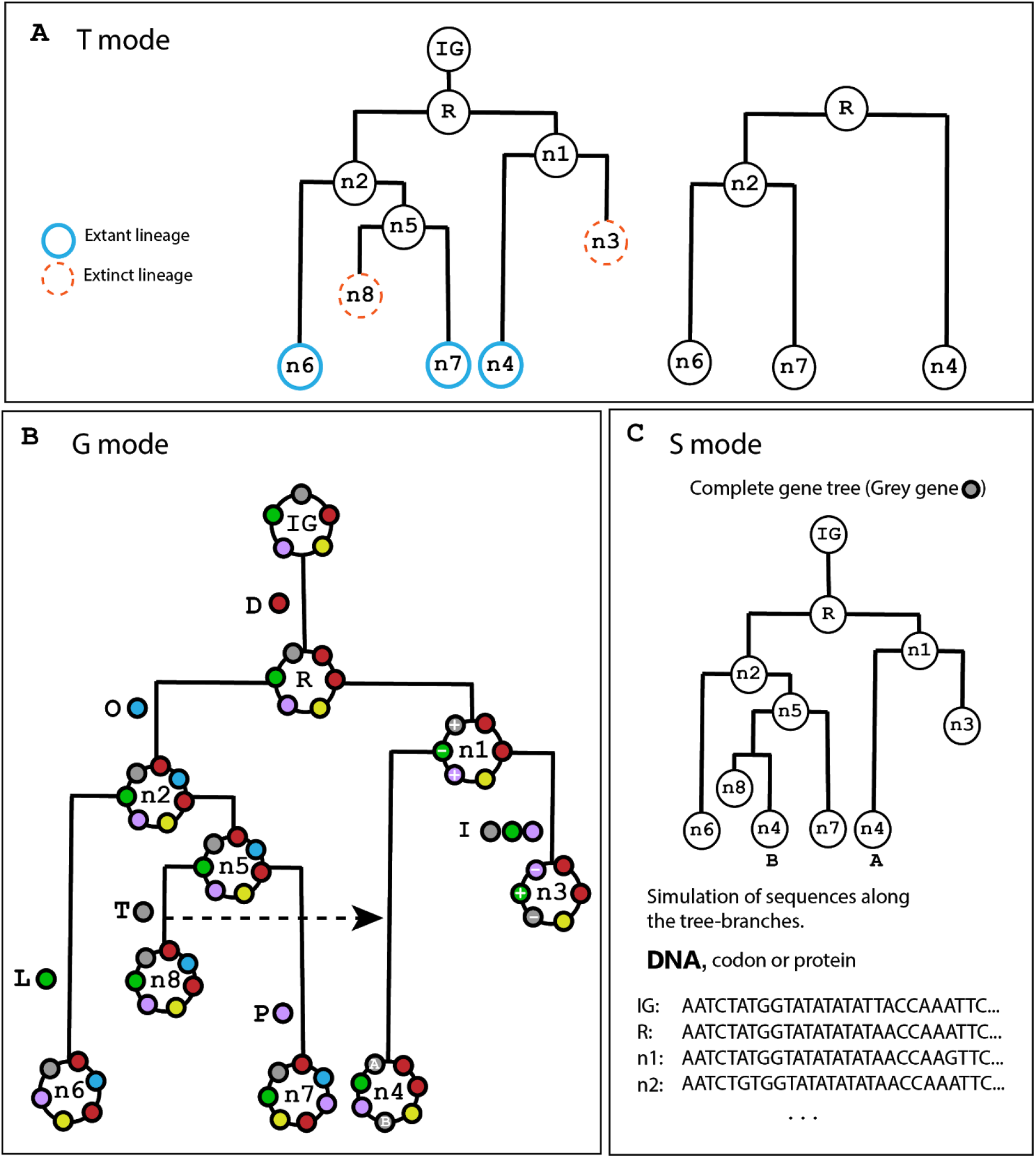
Overview of the three steps of the Zombi simulator. **A**: In T mode, Zombi simulates a species tree using a birth-death process and outputs the pruned version of it by removing extinct lineages. In this example, lineages n3 and n8 go extinct before the simulation ends. **B**: in **G** mode, a circular genome evolves along the branches of the complete species tree obtained with the **T** mode by Duplications (D), Originations (O), Inversions (I), Transpositions (P), Losses (L) and Transfers (T) of genes. The simulation starts with the initial genome (IG) containing a number of genes determined by the user. Each gene has an orientation (+ or −) that is determined randomly and represents the direction of the gene in the coding strand. Here, the IG is composed of five genes (small coloured circles). Various events affecting different genes and their impact on the genome structure are indicated along the branches. The inversion events not only modify the positions of the genes but also change their orientation. **C**: in **S** mode, Zombi can be used to simulate codon, nucleotides and amino acids along the branches of the gene family trees. Here, the gene tree of the grey coloured gene family from **B** has been depicted.

The **T** mode simulates a species tree using a forward simulation under the birth-death model (Kendall 1948), using the Gillespie algorithm (Gillespie 1977), which is the standard method for simulating arbitrarily complex continuous time Markov processes (Figure S1). While more efficient and accurate methods exist to simulate the reconstructed tree, (see e.g. (Hartmann, Wong, and Stadler 2010)), taking into consideration unrepresented (extinct and unsampled) species requires simulating the complete species tree, which includes all extinct and unsampled branches of the phylogeny (Szöllősi et al. 2013). This tree is subsequently pruned to obtain the reconstructed tree, by removing all the lineages that did not survive until the end of the simulation (Figure 1A).

The **G** mode simulates the evolution of genomes along the branches of the complete species tree (Figure 1B), using also the Gillespie algorithm (Figure S2) to account for six possible genome-level events: duplications, losses, inversions, transpositions, transfers and originations. Each of the first five events are characterized by two parameters: the first one is the effective rate, that controls the frequency and fixation probability; the second one controls the extension, i.e., the number of contiguous genes simultaneously affected by the event. Originations of new genes occurs one by one and therefore only a single effective rate parameter is needed. Once the simulation reaches the end, Zombi outputs a list containing each event that has occured in the simulation for every gene family (all genes that share a common origin). Besides, the gene trees of each family are reconstructed by combining both species-level events (Speciations and Extinctions) and genome-level events (Duplications, Transfers and Losses). When a Transfer event occurs, the recipient lineage is randomly chosen from all the lineages alive at that time. The user can make the frequence of transfers to be higher between closely related lineages (Ochman, Lawrence, and Groisman 2000) (Figure S3). Inversions and transpositions do not modify the topology of the tree but add an extra layer of complexity by changing the neighborhood of genes, which is especially relevant when genome-level events affect more than one gene at a time (Figure S4). The gene family trees are also pruned to present the user the trees that can be expected to recover from most real-data analyses, removing all extinct lineages and gene branches that do not arrive until the present time.

The **S** mode, finally, simulates gene sequences (at either the codon, nucleotide or protein level) along the gene family trees (Figure 1C). The user can modify the scaling of the tree to better control the number of substitutions that take place per unit of time, and thus simulate fast or slow-evolving genes.

## 3. Advanced features

In addition to the basic features presented above, “advanced” modes of Zombi (listed in Table S2) can be used to obtain richer and more realistic evolutionary scenarios. For example, it is possible to use a species tree input by the user, to generate species trees with variable extinction and speciation rates, or to control the number of living lineages at each unit of time (Figure S5). At the genome level, Zombi can simulate genomes using branch-specific rates (Gu mode, allowing the user to simulate very specific scenarios such as one in which a certain lineage experiences a massive loss of genes), gene-family specific rates (Gm mode, which makes easier the process of using rates estimated from real datasets) and genomes accounting for intergenic regions (Gf mode) of variable length (drawn from a flat Dirichlet distribution (Biller et al. 2016). At the sequence level, finally, the user can fine-tune the substitution rates to make them branch specific.

Zombi provides the user with a clear and detailed output of the complete evolutionary process simulated, including, the reconciled gene trees with the species tree in the RecPhyloXML reconciliation standard (Duchemin et al. 2018).

## 4. Performance and validation

Simulations with Zombi are fast: with a starting genome of 500 genes and a species tree of 2000 taxa (extinct + extant), it takes around 1 minute on a 3.4Ghz laptop to simulate all the genomes (Figure S6).

We validated that the distribution of waiting times between successive events was following an exponential distribution (Figure S7 and S8), that the distribution of intergene sizes at equilibrium was following a flat Dirichlet distribution, as expected from Biller et al. 2016 (Figure S9), that the number of events and their extension occurs with a frequency according to their respective rates (Figure S10) and that the gene family size distribution followed a power-law when duplication rates are higher than loss rates and stretched-exponential in the opposite case (Szollosi and Daubin 2011; Reed and Hughes 2003) (Figure S11). We also checked by hand the validity of many simple scenarios to detect possible inconsistencies in the algorithm.

## 5. Implementation

Zombi is implemented in Python 3.6. It relies on the ETE 3 toolkit (Huerta-Cepas, Serra, and Bork 2016) and the Pyvolve package (Spielman and Wilke 2015). It is freely available at https://github.com/AADavin/ZOMBI along with detailed documentation and two tutorials in a wiki page.

## Acknowledgments

A.A.D. and G.J.Sz. received funding from the European Research Council under the European Union’s Horizon 2020 research and innovation programme under grant agreement no. 714774., in addition, G.J.Sz. was supported by the grant GINOP-2.3.2.-15-2016-00057. T.T and D.M.d.V received funding from grant ANR-18-CE02-0007-01 (“STHORIZ”). We thank Vincent Daubin, Wandrille Duchemin, Nicolas Lartillot and Thibault Latrille for insightful discussions during the preparation of this manuscript.

## Supplementary Materials

**Table S1:**
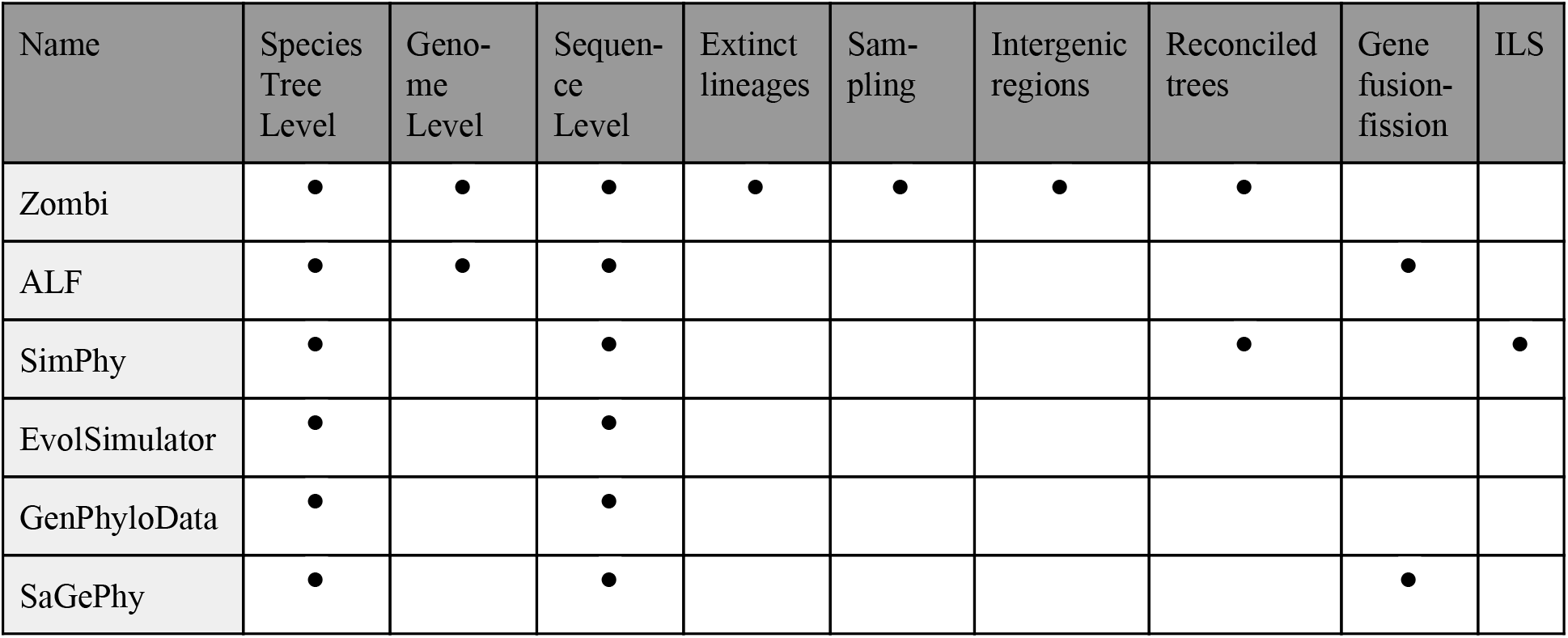
Comparison of the features available in the main evolution simulators. Zombi (this paper), ALF (Dalquen et al. 2011), SimPhy (Mallo, De Oliveira Martins, and Posada 2016), EvolSimulator (Beiko and Charlebois 2007), GenPhyloData (Sjöstrand et al. 2013) and SaGePhy (Kundu and Bansal 2019). The features presented are whether the tool is capable of simulating species trees (Species Tree level), genomes (Genome level, meaning that it considers the structure of the genome, i.e. the physical adjacencies of genes in a genome), sequences (Sequences level), the presence of extinct lineages (Extinct lineages), the possibility of sampling species integrated in the simulator and pruning gene trees according to the species sampled (Sampling), the simulation of intergenic regions (Intergenic regions), outputting reconciled trees (Reconciled trees), considering fusion and fission of genes (Fusion-fission of genes) and producing ILS-induced gene tree/species tree discrepancy (ILS).

**Table S2.**
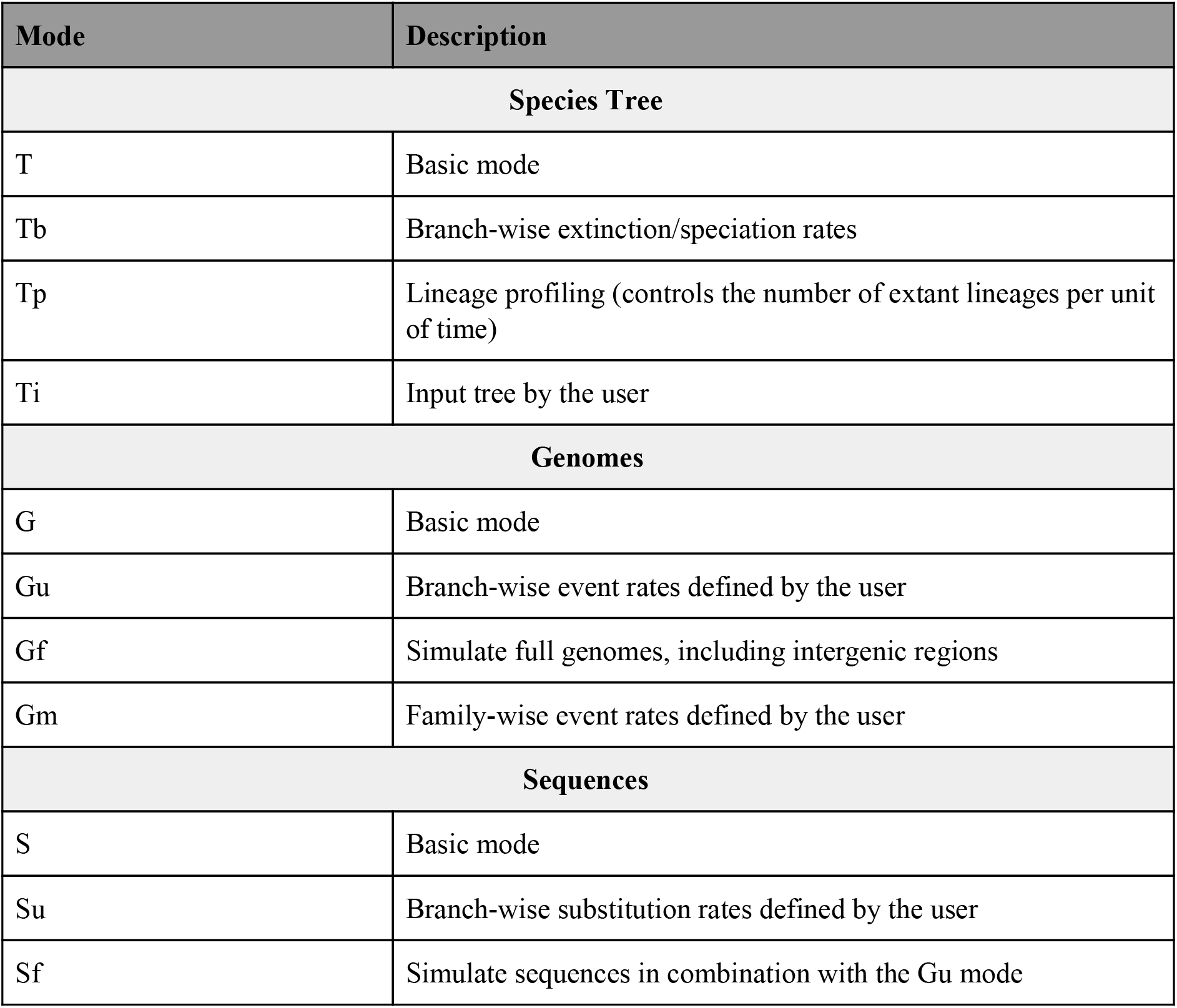
Zombi modes. Zombi implements a total of 11 different modes assigned to three main categories (Species Tree, Genome and Sequence). The basic mode of each category is explained in the main text of this paper.

**Figure S1.**
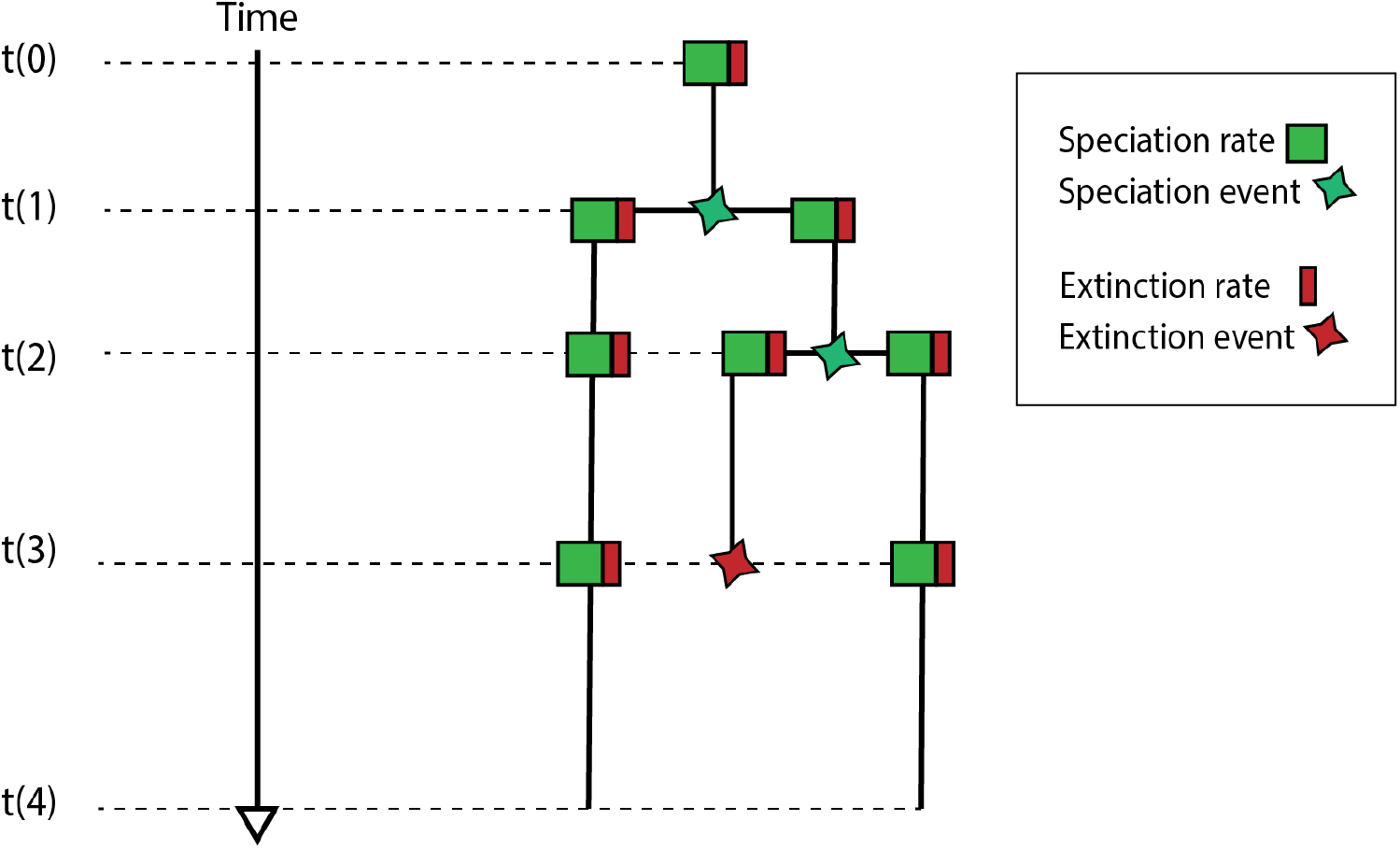
The Gillespie algorithm at the Species Tree level. At these level, there are two possible events: Speciations and Extinctions. Zombi starts with a single lineage at t = 0. To compute the time of occurrence of the next event, a number (t’) is sampled from an exponential distribution with a rate equals to the sum of the rates of the individual event times the number of active branches. Then, a branch is chosen at random from all active branches, and the specific event that it undergoes is chosen according to their relative weights. All the active lineages increase their branch length in t’ units and a new t’ is computed repeating the same procedure. The number of active lineages increases by 1 in the case of a Speciation event or decreases by one in the case of an Extinction. The simulation stops until the total number of lineages reaches a number chosen by the user or the total length of the tree from the initial position to the active leaves attains a certain distance, also controlled by the user. To avoid dead lineages and speciation at the very end of the simulation, the last step of the simulation only increases the branch-length of all the active lineages but does not introduce a new event.

**Figure S2.**
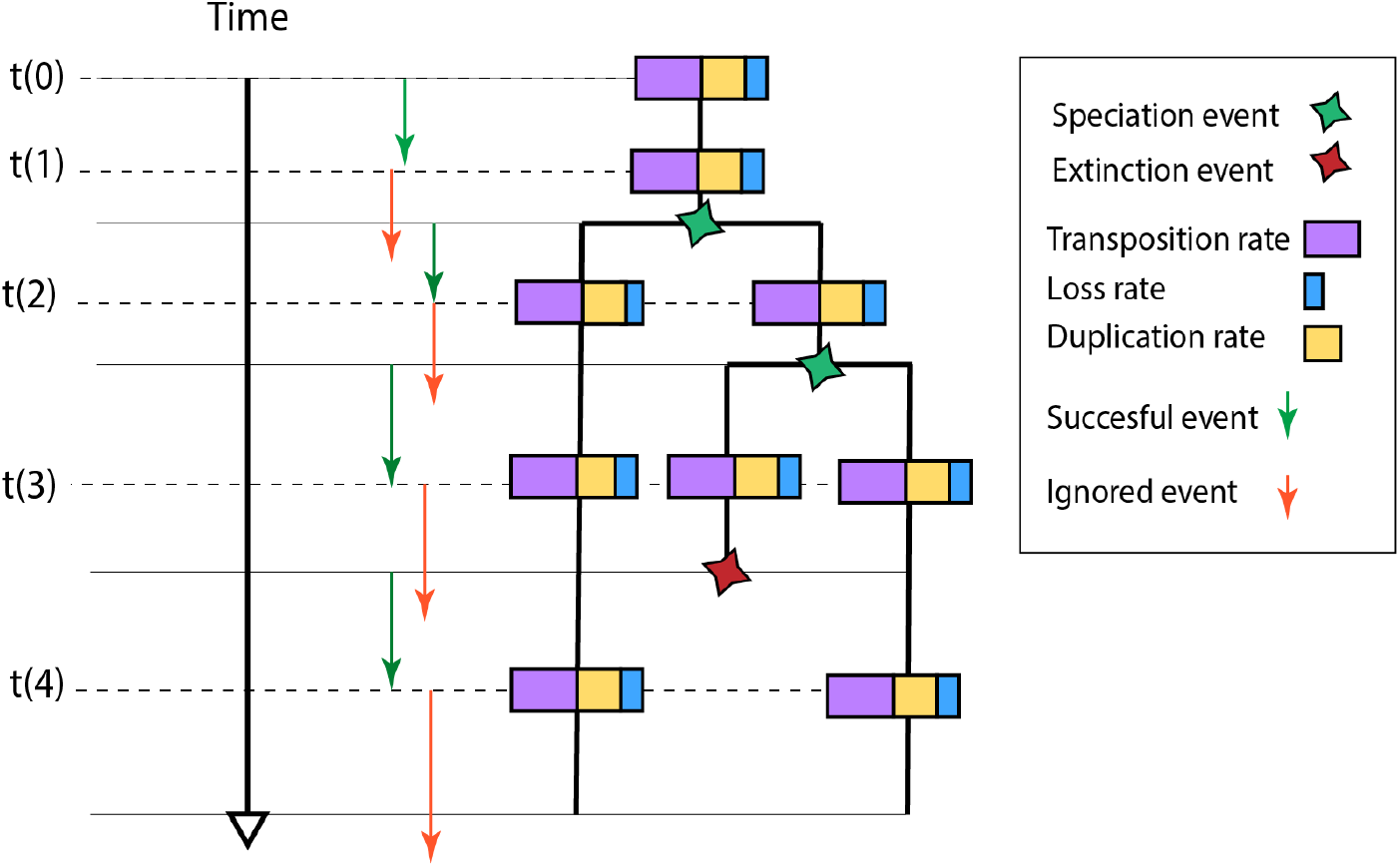
The Gillespie algorithm at the Genome level. A different set of events is used to model the Genome evolution (Duplications, Transfers, Losses, Transpositions, Inversion and Originations). In this example, rates of Origination, Inversion and Transfers are set to 0. The simulation starts at t(0), when the occurrence of the next event is determined by sampling from an exponential distribution with a parameter equal to the sum of all the rates of the active lineages (represented by the squared colours). The underlying patter of speciations and extinctions of the Species Tree is taken into account. If the number sampled form the exponential distribution is smaller than the time remaining for the next Species Tree level event, the event is considered successful and the genome affected is chosen randomly from all active lineages. Then, the specific event taking place is determined according to its rate, as well as its extension, and the affected genes are chosen randomly from all possible contigous positions in the genome. If the event is not successful, it is simply ignored. When a Speciation occurs (determined by the structure of the Species Tree), two identical genomes are created and they continue to evolve independently along the descending branch. Extinction event inactivates the genome evolving within that branch.

**Figure S3.**
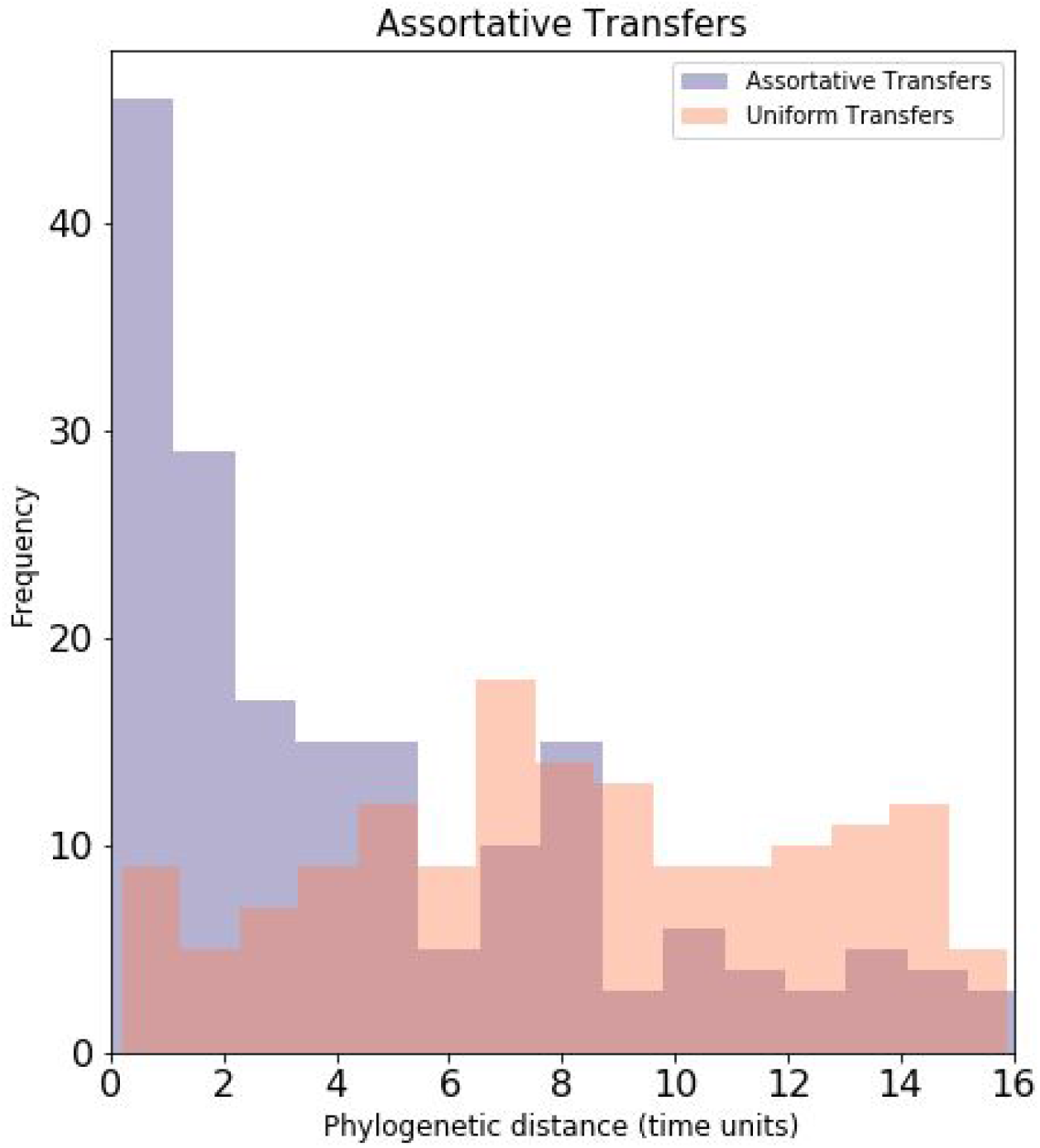
Assortative transfers. By default, when a transfer event occurs, it takes place between two randomly sampled lineages. The user can activate the function assortative transfer, which makes the transfer between two lineages to occur with a probability = *e*^−αδ^ (being α a parameter to control for the strength of the effect and δ the normalized phylogenetic distance). The δ between two nodes is defined as the distance (in time units) between each of the nodes to their common ancestor. In this example, we simulate two datasets in the same Species Tree (30 species). The parameter α was set to 100. We can see how the assortative model of transfer makes transfers between closely related lineages more frequent than the uniform model, a phenomenon that has been observed in real data (Ochman et al. 2000).

**Figure S4.**
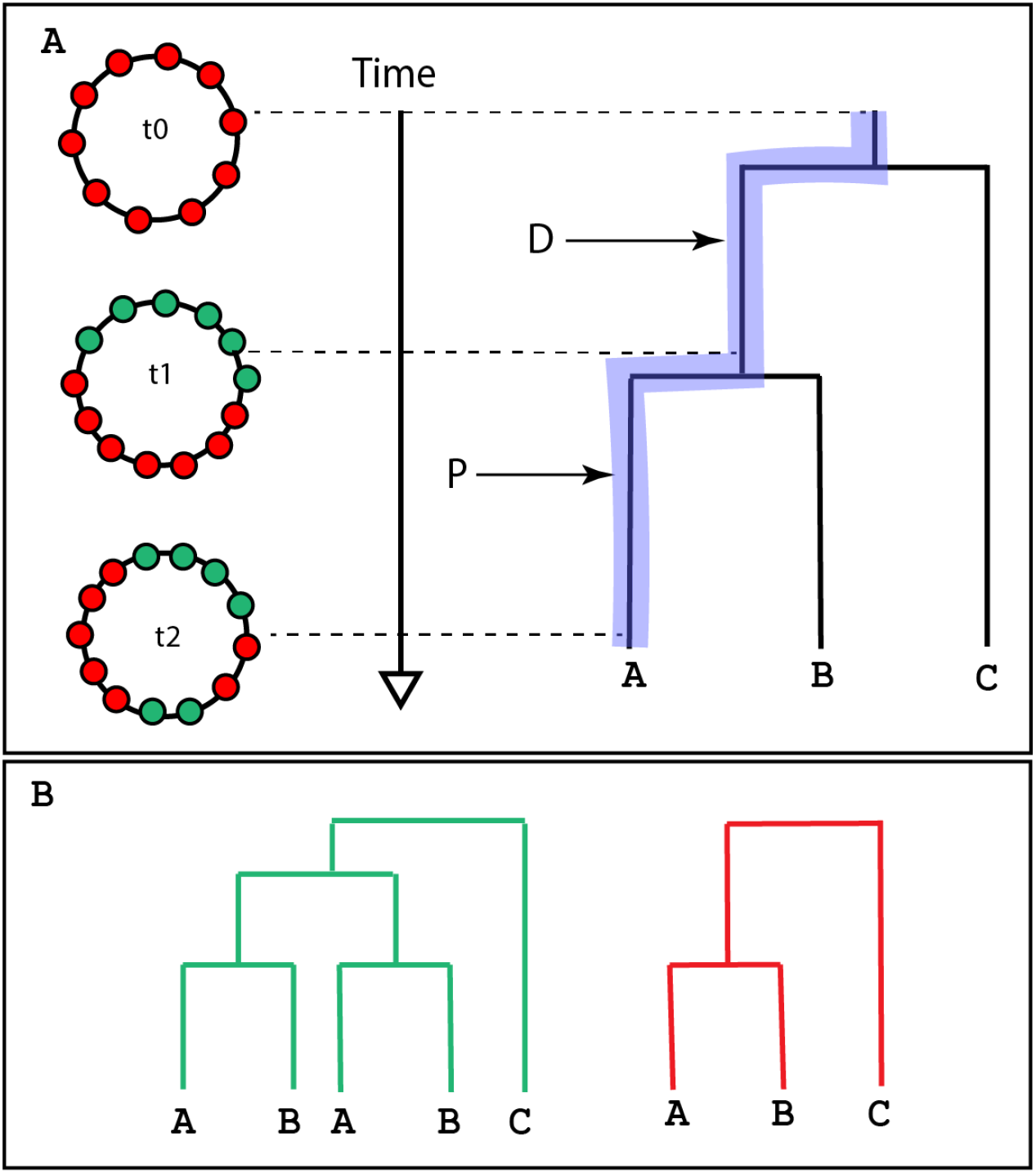
Gene position in Zombi. Simulators like SimPhy (Mallo, De Oliveira Martins, and Posada 2016) consider that all gene families evolve independently one from another. In **Zombi** a single event can affect more than one gene simultaneously. In A we have three snapshots of the genome evolving in the Species Tree on the right, along the blue branches. At time 0, none of the genes has undergone any event. At time 1, the green genes have undergone a duplication. At time 2, some red genes have been transposed within the duplicated genes. In B, the resulting gene trees from this scenario. We can see that in the resulting genomes the genes that, due to the transposition event, share the same duplication event, are shuffled with those that do not. Although inversions and transposition do not alter the topology of the gene trees directly, they can have a big impact when the different events affect more than one gene at a time.

**Figure S5.**
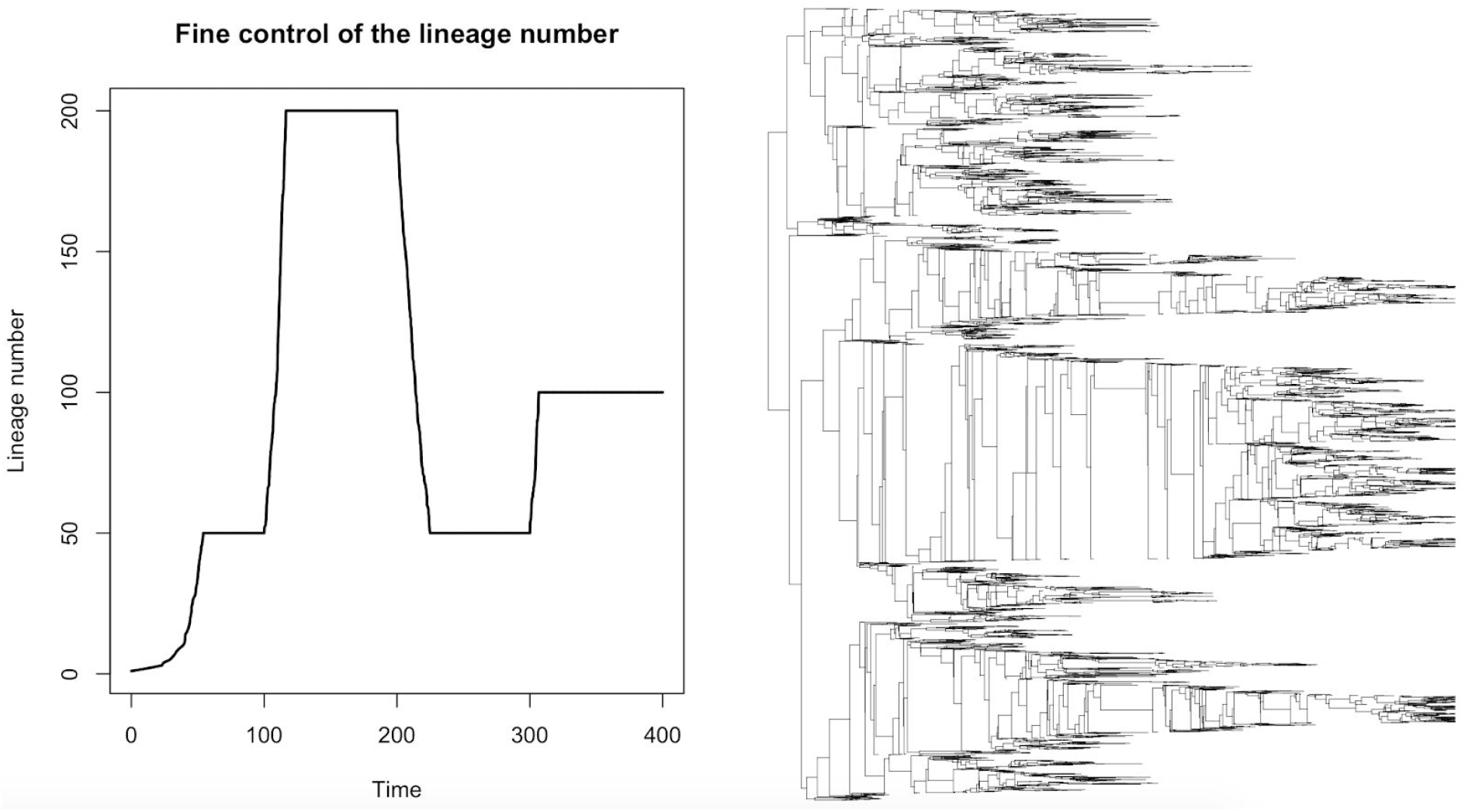
Fine control of the lineage number. Zombi can compute species tree using as input a list of times and the corresponding lineage number that should be attained by that time (in the example t 100 = 50; t 200 = 200; t 300 = 50; t 400 = 100). Zombi tries to attain the lineage number specified for each time interval using the speciation and extinction rates input by the user. At first, there is 1 living lineage and only speciations take place until the number of lineages = 50, number attained in this example when t~50. After that, and because time < 100, the number of lineages reaches an equilibrium in which there is a turnover of species controlled by a parameter also input by the user. Each time that a turnover event takes place two species are randomly sampled in the phylogeny. The first species undergoes a speciation and the second one dies, thus maintaining the total lineage number. The simulation continues until time = 400. In the right panel we can find the resulting species tree.

**Figure S6.**
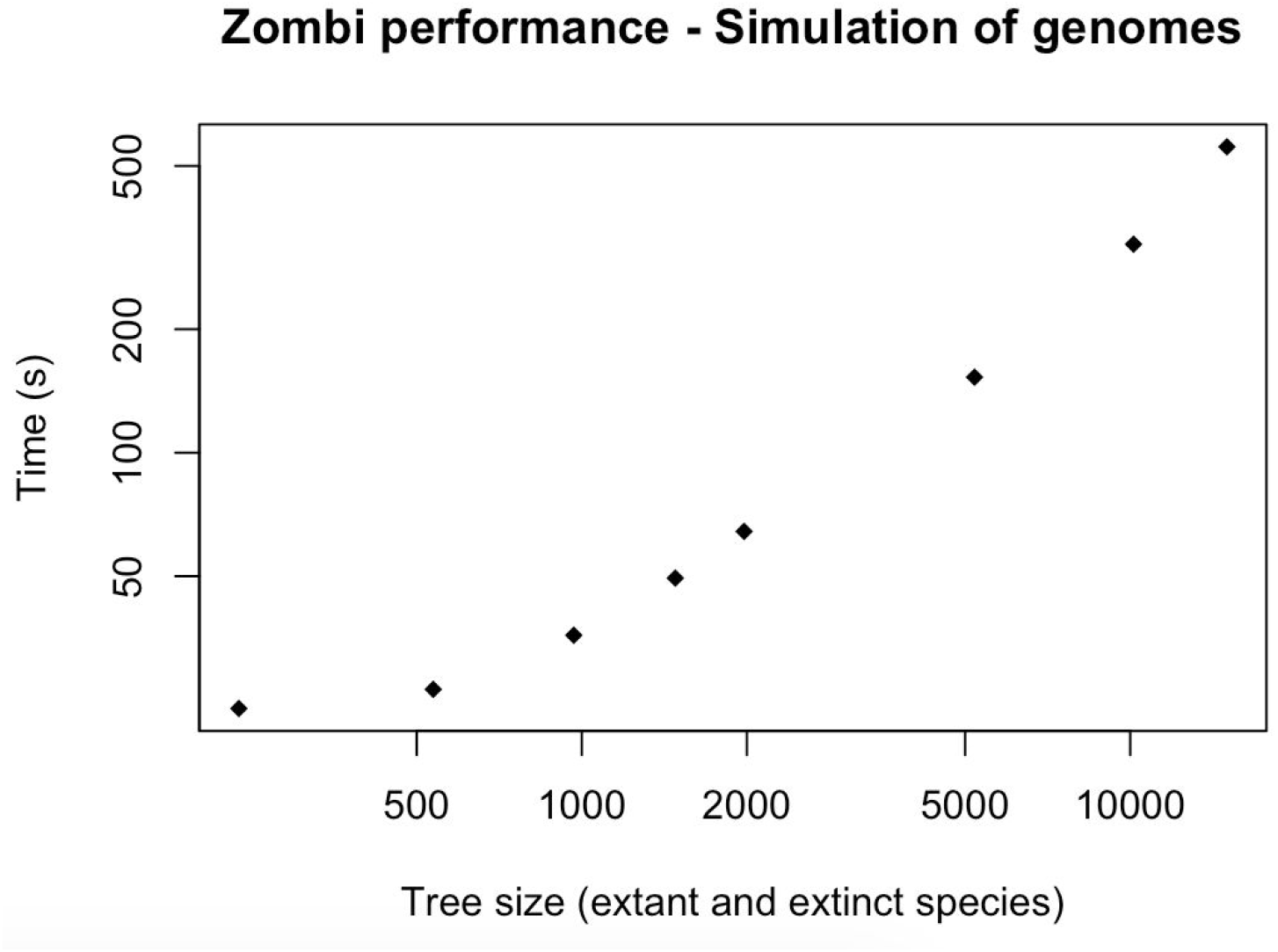
Computing time for different simulation in a computer with a 3,4 GHz Intel Core i5 processor. The rates used were Duplication rate: 0.2, Transfer rate: 0.2, Loss rate: 0.6, Origination rate:0.05, Inversion rate: 0.2, Translocation rate: 0.2. The initial genome was composed of 500 genes. All extension rates were set to 1. Species trees were obtained using by setting Speciation rate: 1 and Extinction rate: 0.5.

**Figure S7.**
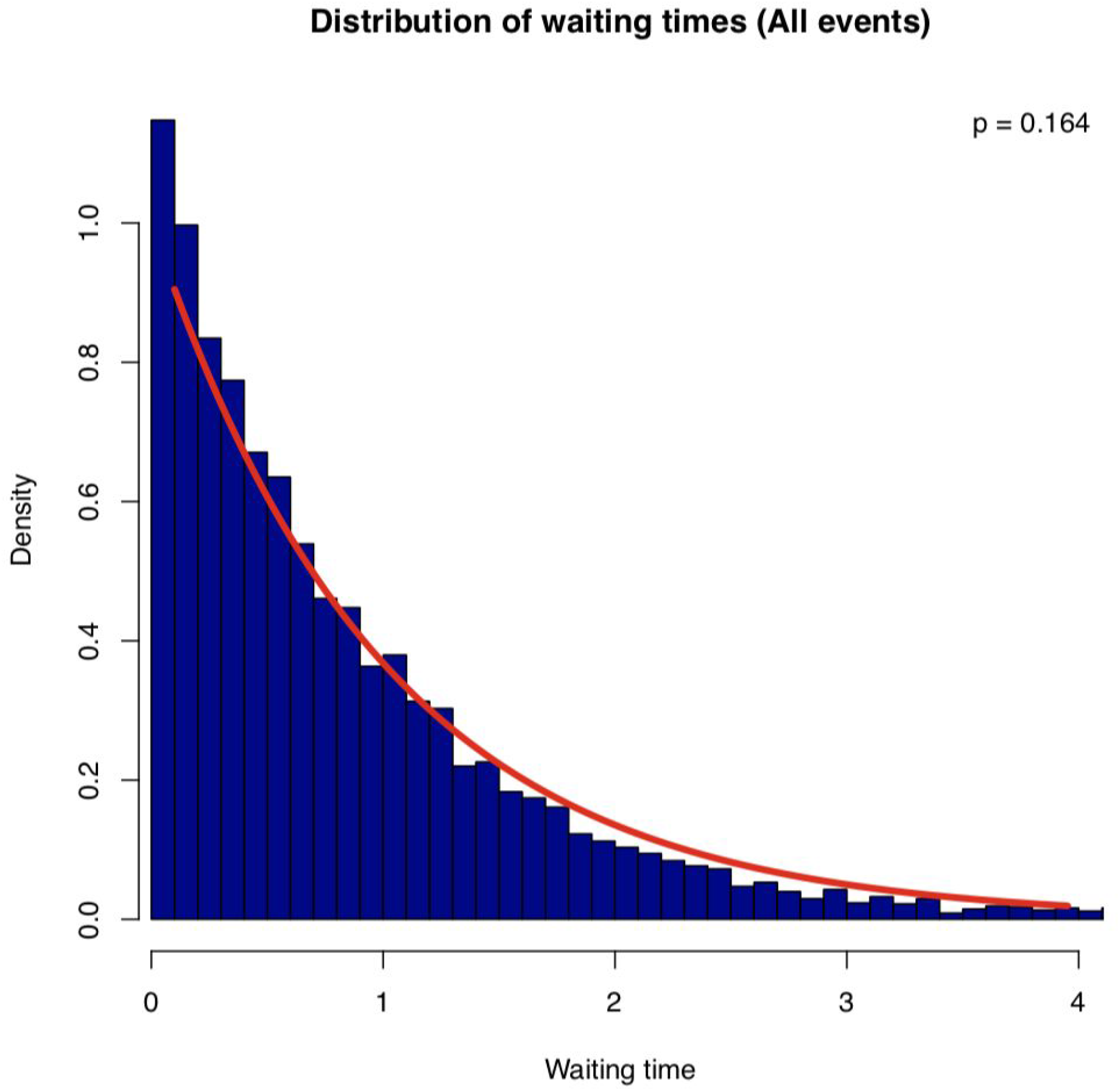
Comparison between the observed distribution of waiting times between consecutive events (duplications, transfers, losses, inversions, translocations and originations, blue bars) and the expected one (red line). The p-value corresponds to a KS test between the empirical distribution and an exponential distribution of the same rate than the same one used in the simulations.

**Figure S8.**
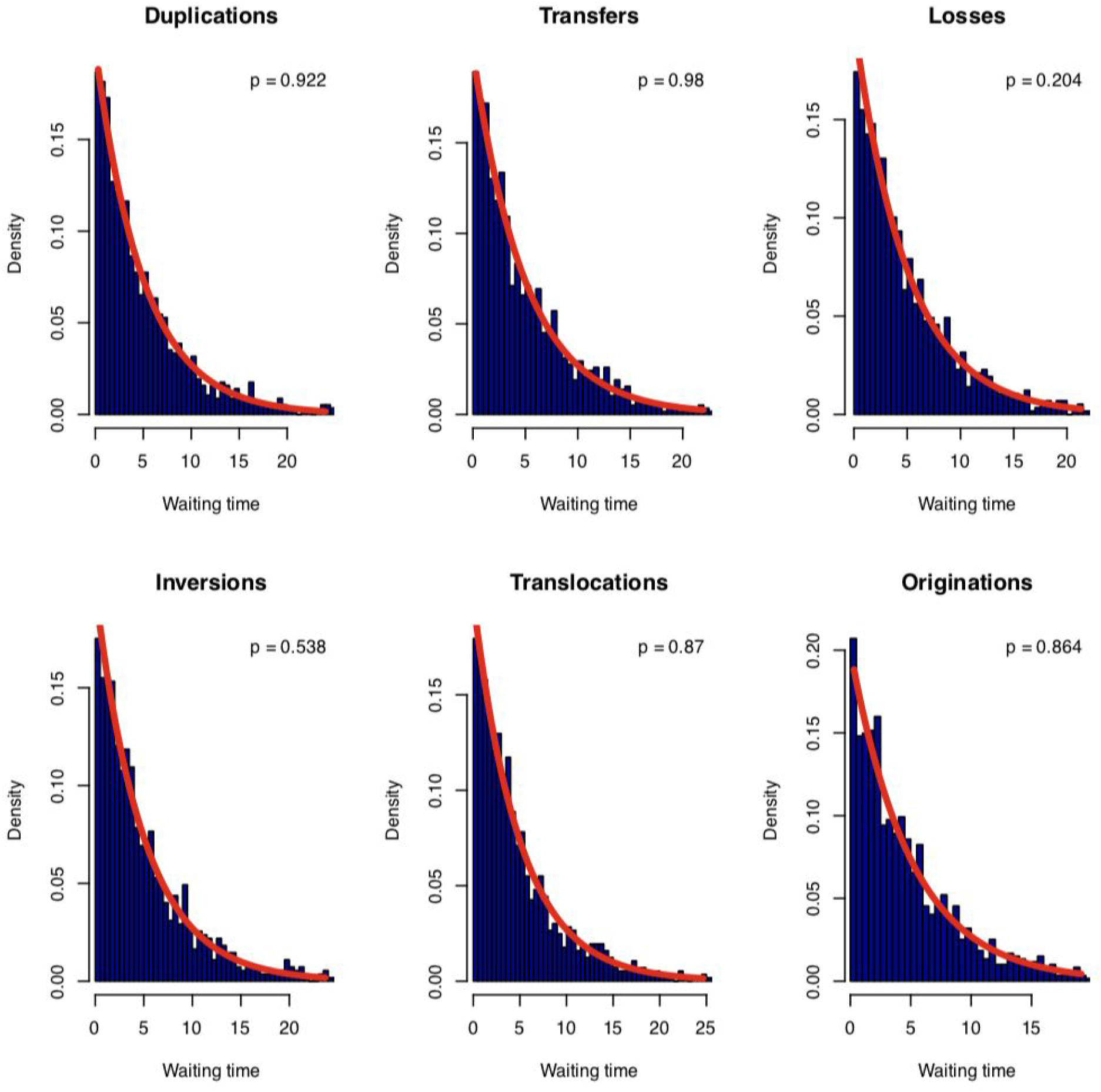
Comparison between the observed (blue bars) and the expected (red line) distribution of waiting times between consecutive events of each type. The p-value corresponds to a KS test between the empirical distribution and an exponential distribution of the same rate.

**Figure S9:**
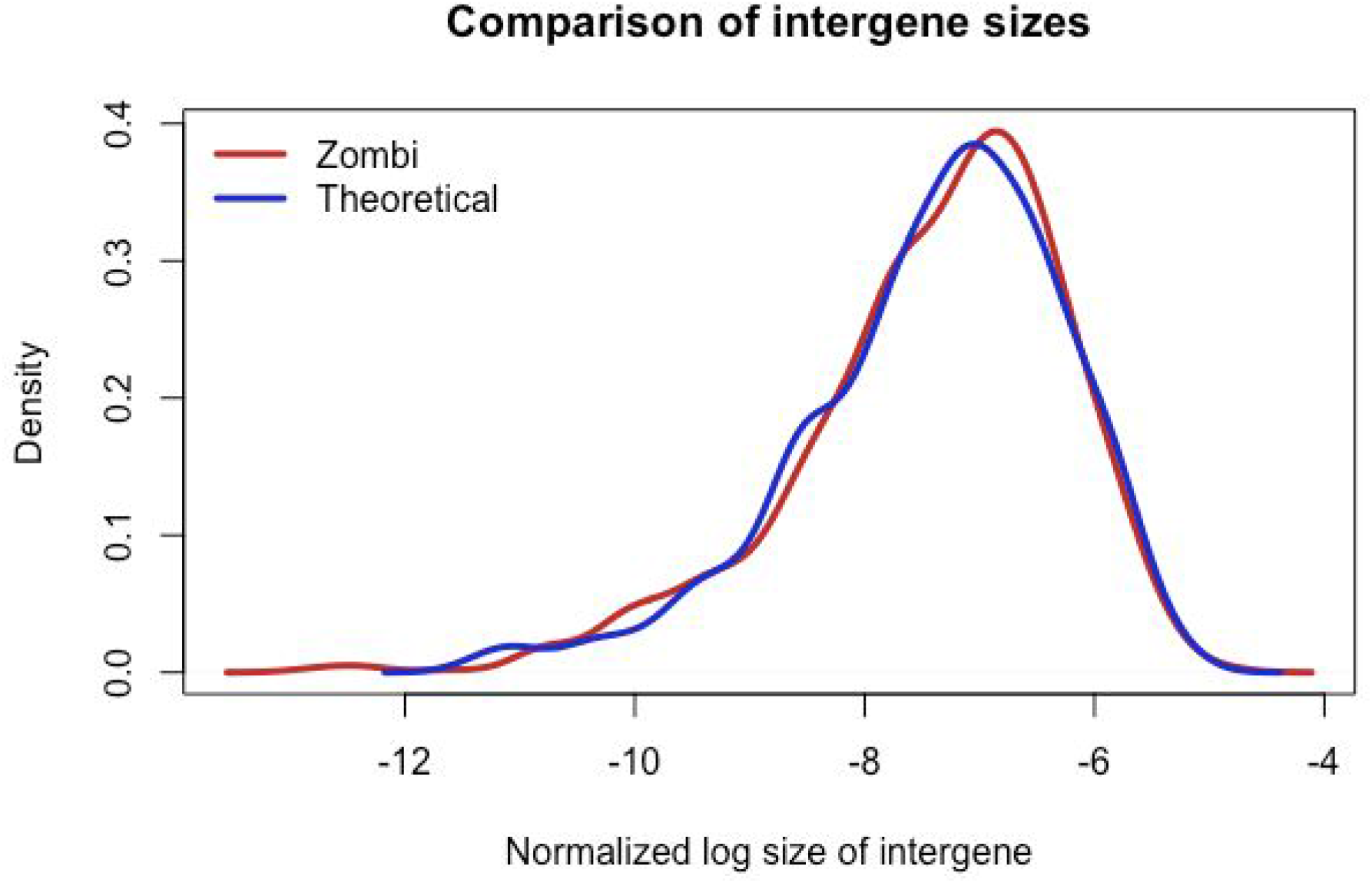
Validation of the mode Gf. To validate the mode **Gf** we simulated a genome with 1000 genes, whose intergene lengths had a constant size of 10000 nucleotides (instead of the Dirichlet as the default option). Then, we made it evolve under many inversion events (~10^6), to see whether at the equilibrium the intergene sizes followed a flat Dirichlet distribution, as expected (see Biller et al. 2016). We compared the obtained values with a randomly generated flat Dirichlet distribution using a KS test and obtained no significant difference (K-S test; p-value = 0.876)

**Figure S10:**
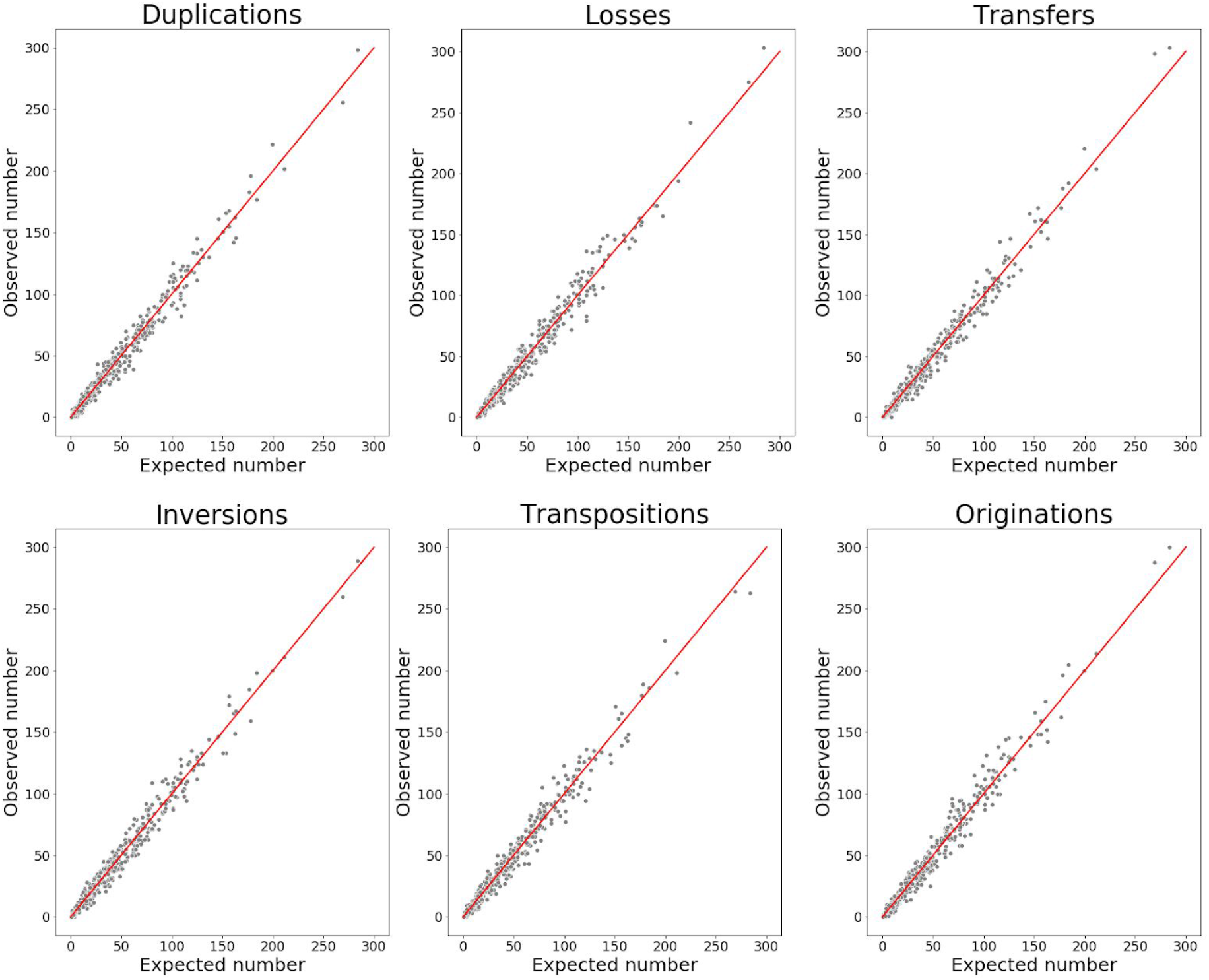
Validation of the rate parameters. We simulate a Species Tree with 100 leaves and the evolution of genomes using the G mode. For every branch of the Species Tree we computed the expected number of events by multiplying the event rate times the branch-length. We plotted the expected number of events against the observed number of events. The red line corresponds to a line with a slope of 1.

**Figure S11:**
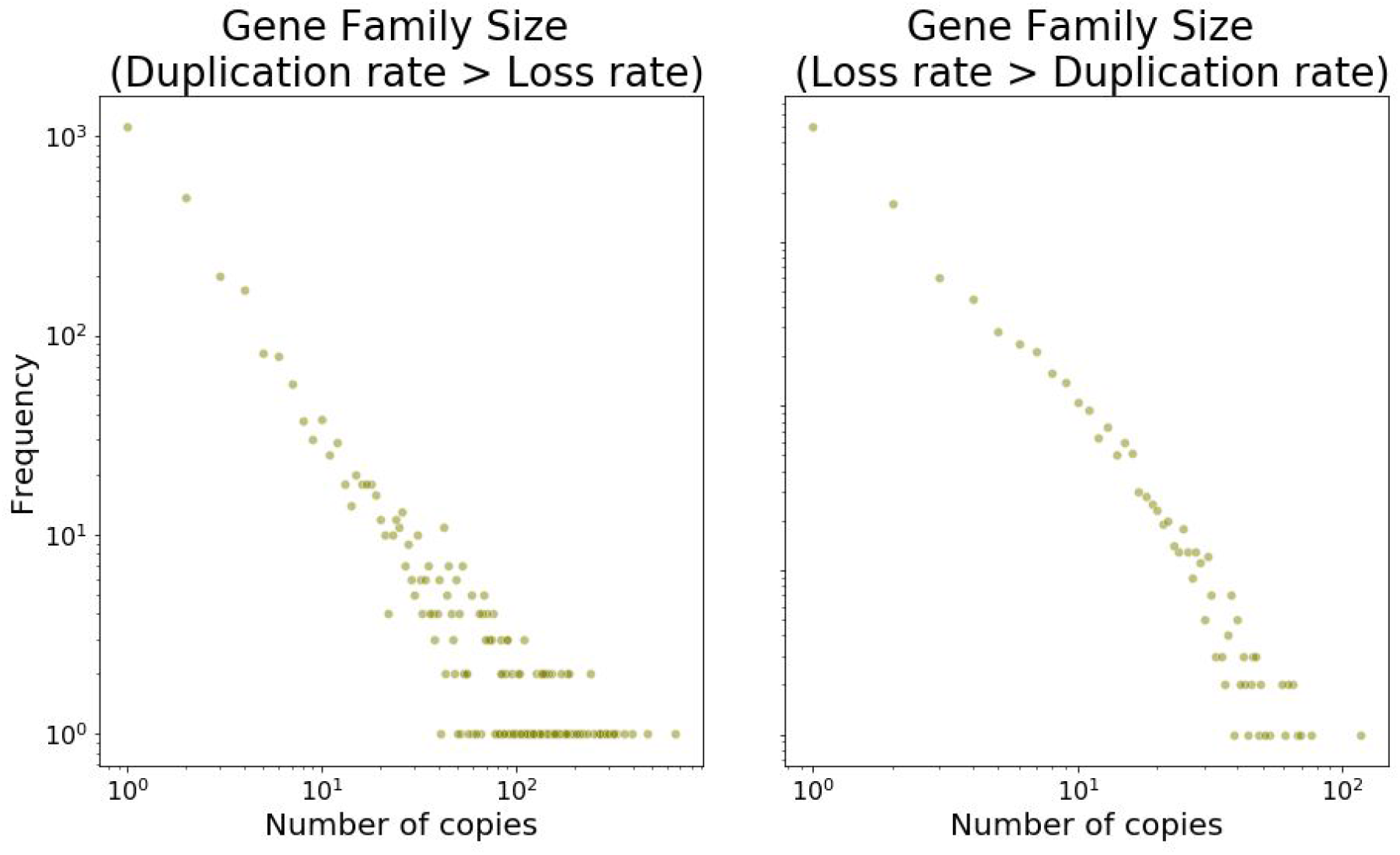
Validation of the mode Gm. In this mode gene families evolve following a birth-death process with specific rates for each family (here we call duplications, D, to births and losses, L, to deaths). It is known that the expected distribution of gene family size follows a power-law distribution when D > L and a stretched exponential if L > D (Szollosi and Daubin 2011; Reed and Hughes 2003). We ran two experiments using the same Species Tree (20 species). In the first one D > L and in the second experiment L > D (the parameter files associated with both experiments can be found in https://github.com/AADavin/ZOMBI/tree/master/Validations. We plotted the frequency against the number of copies for all families with an extant representative in the leaves in log-log axes to inspect visually the distributions and we compared the goodness the support using the Python package Powerlaw (Alstott, Bullmore, and Plenz 2014). We find a clear support for the power-law exponential in the first case (p ~ 3.8^−10) and for the streched-exponential in the second case (p ~5.59 ^ −14)

